# Admix-kit: An Integrated Toolkit and Pipeline for Genetic Analyses of Admixed Populations

**DOI:** 10.1101/2023.09.30.560263

**Authors:** Kangcheng Hou, Stephanie Gogarten, Joohyun Kim, Xing Hua, Julie-Alexia Dias, Quan Sun, Ying Wang, Taotao Tan, Polygenic Risk Methods in Diverse Populations (PRIMED) Consortium Methods Working Group, Elizabeth G. Atkinson, Alicia Martin, Jonathan Shortt, Jibril Hirbo, Yun Li, Bogdan Pasaniuc, Haoyu Zhang

**Author notes:** Address correspondence to: Kangcheng Hou, Bogdan Pasaniuc, Haoyu Zhang.

## Abstract

**Summary:** Admixed populations, with their unique and diverse genetic backgrounds, are often underrepresented in genetic studies. This oversight not only limits our understanding but also exacerbates existing health disparities. One major barrier has been the lack of efficient tools tailored for the special challenges of genetic study of admixed populations. Here, we present admix-kit, an integrated toolkit and pipeline for genetic analyses of admixed populations. Admix-kit implements a suite of methods to facilitate genotype and phenotype simulation, association testing, genetic architecture inference, and polygenic scoring in admixed populations.

**Availability and implementation:** Admix-kit package is open-source and available at https://github.com/KangchengHou/admix-kit. Additionally, users can use the pipeline designed for admixed genotype simulation available at https://github.com/UW-GAC/admix-kit_workflow.

## Introduction

Admixed individuals inherit a mosaic of ancestry segments originating from multiple continental ancestral populations, leading to their complex and diverse genetic backgrounds encompassing a wide spectrum of human genetic variation (Seldin *et al.*, 2011). An understanding of such genetic ancestry mosaicism within admixed populations offers opportunities to gain insights into the origins and health implications of various genetic traits and diseases, contributing to a more comprehensive understanding of human genetics (Wojcik *et al.*, 2019; Tan and Atkinson, 2023).

Despite the genetic richness and crucial insights they can offer, admixed populations remain significantly underrepresented in current genetic studies (Mills and Rahal, 2020). This underrepresentation can be attributed to various challenges, including the complexity of analyzing diverse genetic backgrounds and the lack of efficient tools and standardized practices for handling the genetic data of admixed populations. This gap not only hinders progress in genetic research but also exacerbate health disparities. For example, findings with datasets from European ancestry groups for genetic risk prediction models can introduce bias to personalized risk prevention strategies (Martin *et al.*, 2019; Ding *et al.*, 2023).

To address these challenges, we introduce admix-kit, an integrated and flexible python toolkit along with workflows developed using Workflow Development Language (WDL), specifically designed for the simulation and analysis of genetic data from admixed populations. We anticipate that our proposed software packages and workflows will help overcome these analytical challenges, enabling the inclusion of admixed individuals in future genetic studies.

## Results

### Computational toolkit for analyzing admixed genotypes

We begin by outlining the data structures and computational tools in admix-kit for analyzing admixed genetic datasets. Both genotype and local ancestry data are organized as two matrices of shape N x M x 2 (N and M denote the number of individuals and SNPs respectively, and ‘2’ denotes the two haplotypes; Figure 1a). Given that storage of these matrices often exceeds memory capacity (large N and/or M), we adopt a chunked array representation, implemented with the Dask python library (Rocklin, 2015). Each chunk is loaded from disk on demand, thus conserving memory by loading data only when needed and facilitating large-scale analyses. We employ the pgenlib Python API (URLs) to read phased genotype. Local ancestry matrices are stored in a compressed format that leverages their contiguous nature (local ancestry for nearby SNPs are often identical within each individual). By translating genotype and local ancestry matrices into local-ancestry-specific (LAS) genotype dosages, we have also implemented a set of utility functions tailored for LAS genetic analysis, including LAS allele frequencies, polygenic scores and phenotype modeling that allow for LAS genetic architecture (Figure 1b).

**Figure 1:**
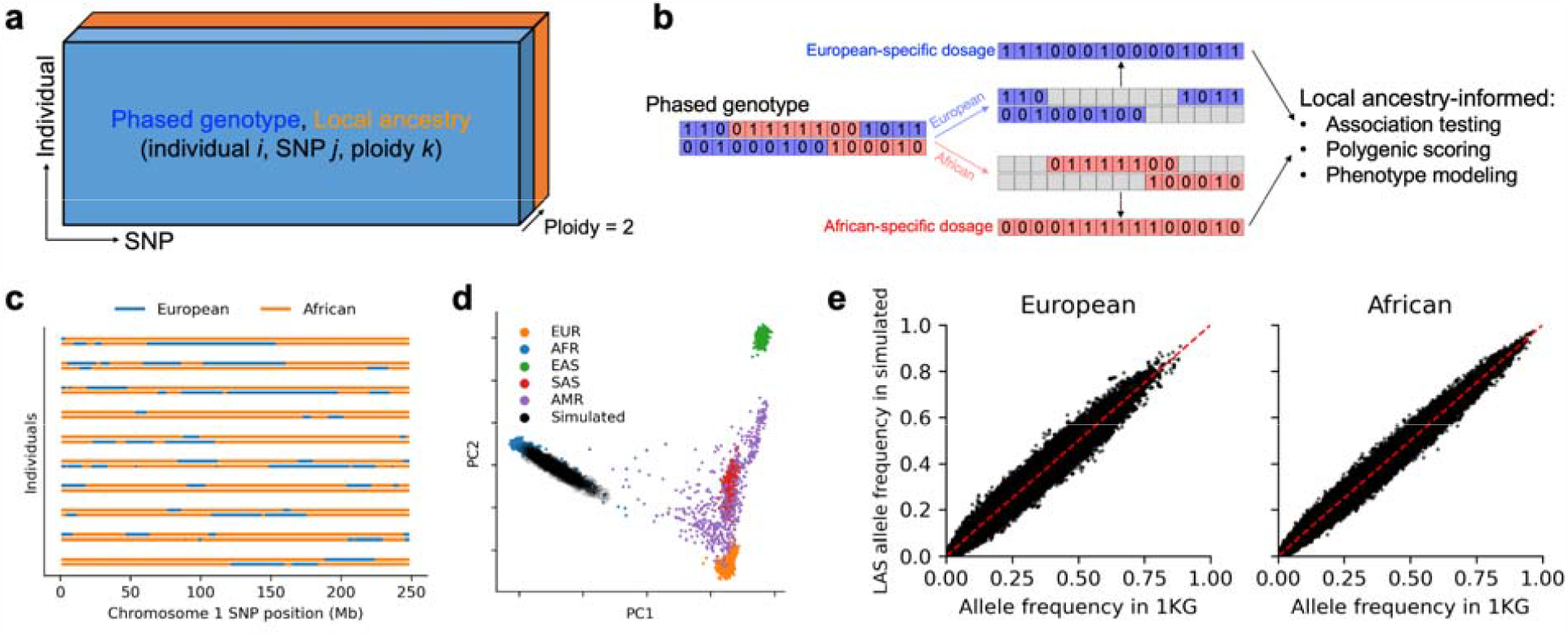
Overview of admix-kit’s data structure, functionality, and illustrative analyses using a simulated dataset (a) Local ancestry and phased genotypes are stored in matrix format. Individual-specific and SNP-specific covariates are stored as two tables with matching orders. (b) Analysis based on ancestry-specific genotype dosage. Starting with a phased genotype for an individual (0/1 denotes presence of minor allele, blue/red color denotes European/African local ancestry), genotypes are separated into ancestry-specific dosages. Local ancestry-informed downstream analyses can be subsequently performed. (c) visualization of local ancestry tracts. (d) Consistency of genome-wide genetic ancestry of simulated dataset (e) Consistency of allele frequencies from the simulated admixed genotypes.

### Workflow for simulating admixture genotypes

Genotype simulation is essential to facilitate testing and benchmarking genetic analysis methodologies. One of the significant challenges lies in simulating admixed genomes, which often becomes the most time-consuming step among common analyses involving admixture. We develop a workflow to specifically address this bottleneck (Figure S1a). We primarily focus on two-way admixture for demonstration while noting our software and pipeline are adaptable to various admixture scenarios. First, we use HAPGEN2 (Su *et al.*, 2011) to expand sets of unique haplotypes within each reference genetic ancestry group (e.g. European/African), preserving the minor allele frequency (MAF) and linkage disequilibrium (LD) structure. Second, using the expanded haplotype sets in both genetic ancestry groups, we simulate the admixture process employing admix-simu (URLs) with parameters for genetic ancestry proportion and the number of admixture generations. This process mimics random mating and recombination events to generate realistic distribution of local ancestry segments, MAF and LD structure for the generated genotypes. To make this simulation process more accessible, we have implemented these functionalities as command-line tools within admix-kit (Figure S1a). In details, `admix hapgen2 --pfile ${src_plink2} --n-indiv ${n_indiv} –out ${expanded_pop}` is used to expand the source population with HAPGEN2. And `admix admix-simu --pfile-list “[‘pop1’, ‘pop2’]” --admix-prop “[0.2,0.8]” -- n-indiv ${n_indiv} --n-gen ${n_gen} --out ${admix}` can be used to simulate the admixture process across source populations. Furthermore, a number of functions are implemented to enable LAS analysis including association testing (Pasaniuc *et al.*, 2011; Atkinson *et al.*, 2021; Hou *et al.*, 2021; Mester *et al.*, 2023) (`admix assoc`), genetic architecture inference (Hou *et al.*, 2023) (`admix genet-cor`) and polygenic scoring (Marnetto *et al.*, 2020; Sun *et al.*, 2022) (`admix calc-partial-pgs`). We also implemented a user-friendly WDL-based workflow for genotype simulation that can be run on cloud-based computing platforms (e.g., AnVIL [https://anvilproject.org/], BioData Catalyst [https://biodatacatalyst.nhlbi.nih.gov/]) (Figure S1b). Users can input essential parameters to define the admixture process and provide the input genotype path of ancestral populations (a set of preprocessed 1,000 genomes dataset is provided for default usage). The workflow will run through each aforementioned step and produce the simulated admixed genotype dataset. The admix-kit software is encapsulated in a publicly available docker image (URLs).

### Example analysis of a simulated dataset

We demonstrate the practicality of admix-kit through analyses of a simulated dataset. All associated code and notebooks have been made publicly accessible (https://github.com/UW-GAC/admix-kit_workflow). This ensures our results are fully reproducible and can be seamlessly deployed in a cloud platform (e.g., AnVIL). We used the AnVIL workflow to simulate N = 1,000 admixed individuals with M = 174K SNPs on chromosome 1 and 2, using a demographic model similar to African American individuals with over 8 generations of admixture and an average ancestry proportion of 80% African and 20% European (Kidd *et al.*, 2012) (ancestry proportion varies by individual). Notably, the genotype simulation took less than 30 minutes with scalability to a much larger number of individuals and SNPs. Using principal component analysis (PCA), we observed that individuals within the simulated dataset are positioned along a cline between individuals labeled as European and African in the 1,000 Genomes reference dataset, suggesting high quality of the simulated genotype dataset (Figure 1c-d). Allele frequencies computed within genotype segments corresponding to the respective local ancestry displayed high consistency with those computed in the reference population, indicating high preservation of MAF structure of the simulated genotype (Figure 1e).

## Discussion

Addressing the underrepresentation of admixed individuals in genetic studies is pivotal not only for scientific necessity but also as a commitment to equity. With this goal in mind, we introduce admix-kit, a comprehensive toolkit and workflow tailored for admixed populations. We anticipate that our software package and workflows will facilitate greater inclusion of admixed individuals in future genetic studies.

Development of software and methodology in genetic studies relies heavily on the use of simulated datasets. These datasets help benchmark performance and facilitate comparisons with existing software. Traditionally, simulated datasets are usually derived from publicly available reference populations. Often, these populations are selected based on a high degree of genetic similarity among individuals in the population (e.g., individuals having all four grandparents from a small geographic region.) For instance, HAPGEN2 has recently been widely used for simulating large-scale genetic datasets that mimic the LD structures of reference populations such as European, African, American, East Asian, and South Asian using data from the 1000 Genomes Project (Su *et al.*, 2011; Ruan *et al.*, 2022; Zhang *et al.*, 2022; Miao *et al.*, 2023). While these simulations can recreate datasets with homogeneous LD as the reference populations, they may not capture the complex population structures seen in admixed populations (Figure S2). Consequently, these sampling conditions are not representative of global human genetic variations. As a remedy, simulating admixture among reference populations can provide datasets that more rigorously test the performance of new software. For example, our simulation pipeline can be used to investigate factors that potentially impact accuracy of ancestry inference (including ancestry composition in reference panel, demographic model of simulated admixed population and error in inferred local ancestry) and to understand how errors in ancestry inference propagate to downstream disease mapping and prediction applications.

Admix-kit holds significant potentials in the development of Polygenic Risk Scores (PRS). The efficacy of PRS is known to hinge on the similarity of the target population to the training population (Ding *et al.*, 2023). With the PRIMED consortium working on methods to improve the performance of PRS in diverse populations, simulations will be pivotal for method evaluation (Kachuri *et al.*, 2023). In this context, we expect that admix-kit will be an essential part of this effort.

## Acknowledgements

This research was funded in part by the National Institutes of Health under awards U01-HG011715 (B.P.). H.Z. was supported by NIH Intramural Research Program. E.G.A. was supported by grant K01MH121659 from the National Institutes of Health/National Institute of Mental Health, the Caroline Wiess Law Fund for Research in Molecular Medicine, and the ARCO Foundation Young Teacher–Investigator Fund at Baylor College of Medicine. Research reported in this publication was supported by the National Institutes of Health for the project “Polygenic Risk Methods in Diverse Populations (PRIMED) Consortium”, with grant funding for Study Sites CAPE (U01HG011715) and FFAIRR-PRS (U01HG011719), and the Coordinating Center (U01HG011697). The content is solely the responsibility of the authors and does not necessarily represent the official views of the National Institutes of Health.

## URLs

Admix-kit: https://github.com/KangchengHou/admix-kit

Admixed genotype simulation workflow: https://github.com/UW-GAC/admix-kit_workflow

Admix-kit docker image: https://dockstore.org/organizations/PRIMED/collections/simulation

Pgenlib: https://github.com/chrchang/plink-ng/blob/master/2.0/Python/pgenlib.pyx

Admix-simu: https://github.com/williamslab/admix-simu

PRIMED consortium: https://primedconsortium.org/

---

**Figure S1:**
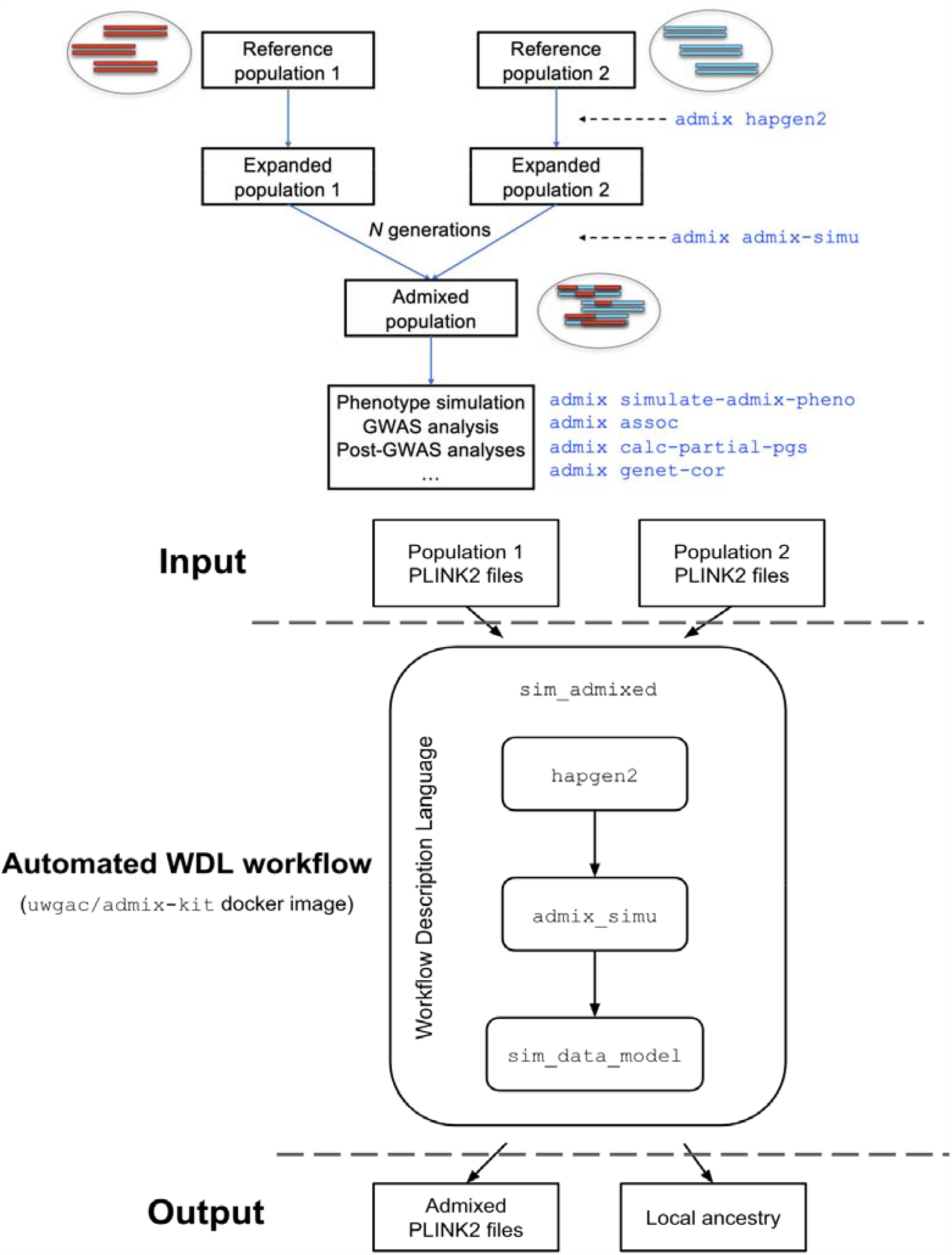
Overview of genotype simulation for admixed populations. (upper panel) Illustration of admixed genotype simulation and the corresponding python API. (lower panel) Illustration of WDL workflow.

**Figure S2:**
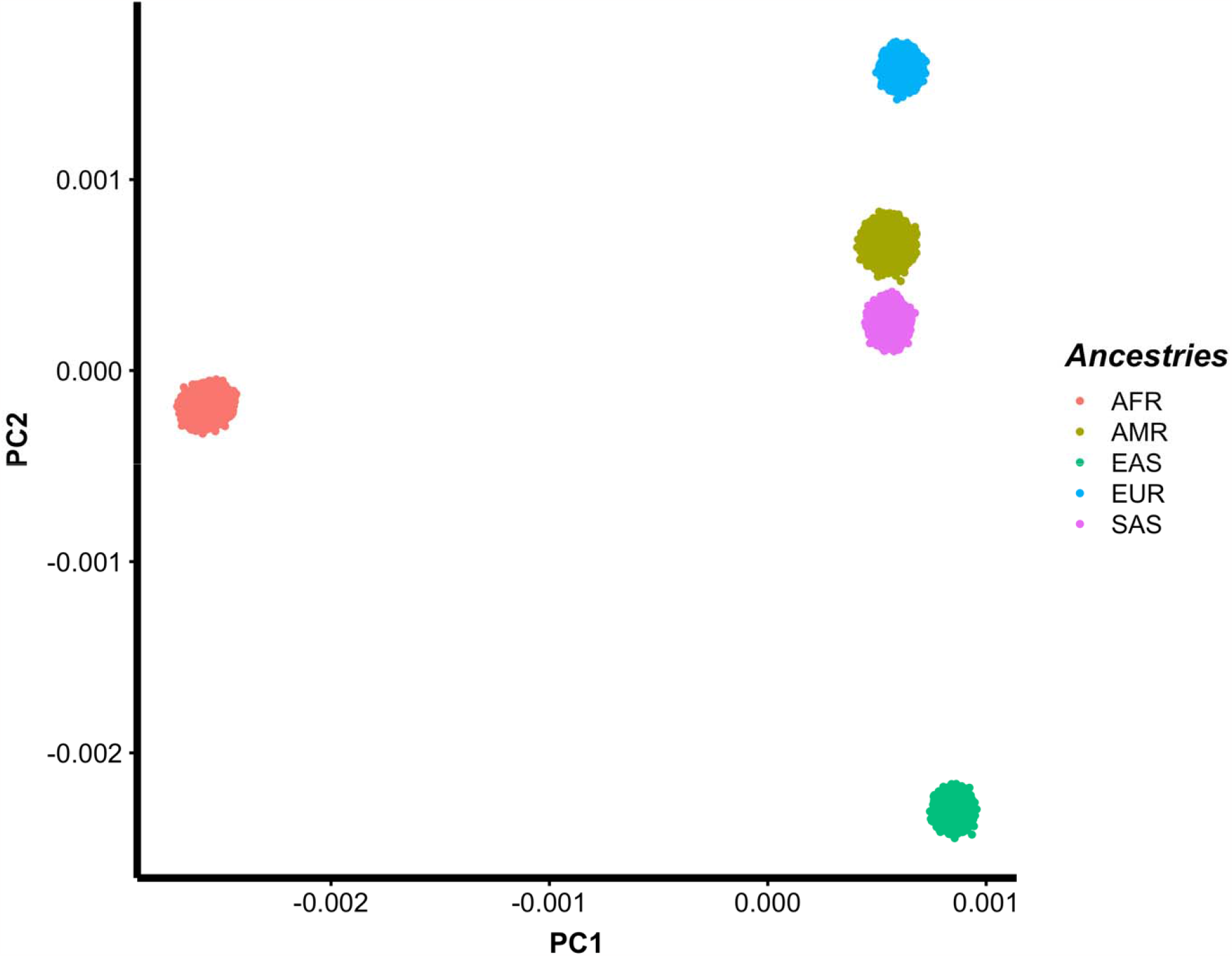
Principal component analysis of 600,000 simulated multi-ancestry subjects using Hapgen2 and 1000 Genomes Project (Phase 3) as references. The 1000 Genomes Project contains 661 African, 347 American, 504 East Asian, 503 European and 489 South Asian subjects. In this simulation, each of the five ancestries contributed 120,000 subjects. The population structure and the continuous genetic ancestry variation in 1000 Genomes Project (observed in Figure 1d) is not observed in these simulated data (e.g., simulated AMR individuals are relatively homogenous).

## References

Atkinson, E.G. et al. (2021) Tractor uses local ancestry to enable the inclusion of admixed individuals in GWAS and to boost power. Nat. Genet., 53, 195–204.

Ding, Y. et al. (2023) Polygenic scoring accuracy varies across the genetic ancestry continuum. Nature, 618, 774–781.

Hou, K. et al. (2023) Causal effects on complex traits are similar for common variants across segments of different continental ancestries within admixed individuals. Nat. Genet., 55, 549–558.

Hou, K. et al. (2021) On powerful GWAS in admixed populations. Nat. Genet., 53, 1631–1633.

Kachuri, L. et al. (2023) Principles and methods for transferring polygenic risk scores across global populations. Nat. Rev. Genet.

Kidd, J.M. et al. (2012) Population genetic inference from personal genome data: impact of ancestry and admixture on human genomic variation. Am. J. Hum. Genet., 91, 660–671.

Marnetto, D. et al. (2020) Ancestry deconvolution and partial polygenic score can improve susceptibility predictions in recently admixed individuals. Nat. Commun., 11, 1628.

Martin, A.R. et al. (2019) Clinical use of current polygenic risk scores may exacerbate health disparities. Nat. Genet., 51, 584–591.

Mester, R. et al. (2023) Impact of cross-ancestry genetic architecture on GWASs in admixed populations. Am. J. Hum. Genet., 110, 927–939.

Miao, J. et al. (2023) Quantifying portable genetic effects and improving cross-ancestry genetic prediction with GWAS summary statistics. Nat. Commun., 14, 1–13.

Mills, M.C. and Rahal, C. (2020) The GWAS Diversity Monitor tracks diversity by disease in real time. Nat. Genet., 52, 242–243.

Pasaniuc, B. et al. (2011) Enhanced statistical tests for GWAS in admixed populations: assessment using African Americans from CARe and a Breast Cancer Consortium. PLoS Genet., 7, e1001371.

Rocklin, M. (2015) Dask: Parallel Computation with Blocked algorithms and Task Scheduling. In, Proceedings of the 14th Python in Science Conference. SciPy.

Ruan, Y. et al. (2022) Improving polygenic prediction in ancestrally diverse populations. Nat. Genet., 54, 573–580.

Seldin, M.F. et al. (2011) New approaches to disease mapping in admixed populations. Nat. Rev. Genet., 12, 523–528.

Su, Z. et al. (2011) HAPGEN2: simulation of multiple disease SNPs. Bioinformatics, 27, 2304–2305.

Sun, Q. et al. (2022) Improving polygenic risk prediction in admixed populations by explicitly modeling ancestral-specific effects via GAUDI. bioRxiv.

Tan, T. and Atkinson, E.G. (2023) Strategies for the genomic analysis of admixed populations. Annu. Rev. Biomed. Data Sci., 6, 105–127.

Wojcik, G.L. et al. (2019) Genetic analyses of diverse populations improves discovery for complex traits. Nature, 570, 514–518.

Zhang, H. et al. (2022) A new method for multi-ancestry polygenic prediction improves performance across diverse populations. bioRxiv, 2022.03.24.485519.

